# Deep sequencing of 16 *Ixodes ricinus* ticks unveils insights into their interactions with endosymbionts

**DOI:** 10.1101/2024.04.22.590557

**Authors:** Paulina M. Lesiczka, Tal Azagi, Aleksandra I. Krawczyk, William T. Scott, Ron P. Dirks, Ladislav Šimo, Gerhard Dobler, Bart Nijsse, Peter J. Schaap, Hein Sprong, Jasper J. Koehorst

**Author notes:** Corresponding author biology. Corresponding author bioinformatics. These authors contributed equally to this work (Paulina M. Lesiczka, Tal Azagi, Hein Sprong, Jasper J. Koehorst).

## Abstract

**Background:** *Ixodes ricinus* ticks act as vectors for numerous pathogens that present substantial health threats. Additionally, they harbour vertically transmitted symbionts, some of which have been linked to diseases. The difficulty of isolating and cultivating these symbionts has hampered our understanding of their biological role, their potential to cause disease, and their modes of transmission. To expand our understanding on the tick symbiont *Midichloria mitochondrii* and on *Rickettsia helvetica*, which has been linked to disease in humans, we utilized deep sequencing on sixteen individual adult female ticks collected from coastal dune and forested areas in the Netherlands.

**Results:** By employing a combination of second and third-generation sequencing techniques, we successfully reconstructed the complete genomes of *M. mitochondrii* from eleven individuals, *R. helvetica* from eight individuals and the mitochondrial genome from all ticks. Additionally, we visualised the location of *R. helvetica* in tick organs and constructed genome-scale metabolic models (GEMs) of both symbionts to study their environmental dependencies.

Our analysis revealed a strong cophylogeny between M. mitochondrii and mitochondrial genomes, suggesting frequent maternal transmission. In contrast, the absence of cophylogeny between R. helvetica and the mitochondrial genomes, coupled with its presence in the receptaculum seminis of I. ricinus females, raises the possibility of paternal transmission of R. helvetica. Notably, the genetic diversity of R. helvetica was found to be very low, except for the rickA virulence gene, where the presence of up to thirteen insertions of a33nt-long repeat led to significant variability. However, this variation could not account for the differences in infection prevalence observed across eight distinct locations in the Netherlands.

**Conclusions:** By employing deep sequencing, it becomes feasible to extract complete genomes and genetic data of symbionts directly from their host organisms. This methodology serves as a robust means to gain fresh insights into their interactions. Our observations, which suggest paternal transmission of *R. helvetica*, a relatively unexplored mode of transmission in ticks, require validation through experimental investigations. The genetic variations identified in the *rick*A virulence gene of *R. helvetica* have the potential to influence the infectivity and transmission dynamics of *R. helvetica*

## 1. Introduction

*Ixodes ricinus*, the most common tick species in the Palearctic realm, carries a variety of facultative symbionts, some of which can cause infectious diseases in humans. The most common tick-borne diseases associated with *I. ricinus* are Lyme disease (LB) which is caused by *Borrelia burgdorferi sensu lato* and tick-borne encephalitis (TBE), which is caused by tick-borne encephalitis virus [1]. In addition, *I. ricinus* transmits several emerging pathogens such as *Rickettsia helvetica*, *Anaplasma phagocytophilum*, *Borrelia miyamotoi*, *Neoehrlichia mikurensis*, and several *Babesia* species [2]. Infections with these pathogens have been associated with human disease in cross-sectional and case studies, but their pathogenic potential has not been fully understood [3, 4]. These microorganisms can influence the tick host’s metabolism, development, reproduction, stress defence, and immunity [5] and may indirectly contribute to the transmission dynamics of tick-borne pathogens, significantly impacting human and animal health [6].

The biological role, virulence, and transmission dynamics of tick symbionts and some of the pathogens are often difficult to study due to challenges in isolating and culturing them. A notable example is *Midichloria mitochondrii*, initially discovered in the ovaries of ticks. *M. mitochondrii* has been found to be highly prevalent in various tissues of *I. ricinus* [7–10]. Since its discovery, *Midichloria* symbionts have been detected in several tick species across different genera, albeit with varying prevalence [11], and are believed to contribute to detoxification, vitamin supply, and energy-related functions, based on their observed interactions and effects within host cells [12]. While *M. mitochondrii* has not been associated with human disease and is only considered in the context of tick-symbiont interactions, other symbionts such as *Rickettsia helvetica* cannot be so clearly defined. *Rickettsia helvetica* is a Gram-negative, obligate intracellular bacterium formerly included in the Spotted Fever Group Rickettsiae (SFGR) [13]. *Rickettsia helvetica* can be transmitted by tick bites and has been associated with human disease. Only a few human cases have been reported in Europe [14–19] and detection in cerebrospinal fluid of suspected Lyme neuroborreliosis patients has been shown in the Netherlands and Denmark. However, it is unclear whether infection with *R. helvetica* can cause disease or might change the course of Lyme disease [20, 21]. Moreover, studies on the involvement of *R. helvetica* in sarcoidosis seem to be contradictory [22]. Thus, the extent of *R. helvetica*’s infectivity and pathogenicity is still under investigation, and little is known about its ecoepidemiology [23, 24]. DNA of *R. helvetica* has only been sporadically detected in wildlife [25–29] with no clear animal reservoir. This symbiont is transmitted vertically from one generation of ticks to the next and has been repeatedly isolated from questing *I. ricinus* in European countries [30], which are currently thought to be the main reservoir for *R. helvetica* in nature [25]. *Rickettsia helvetica* infection rates in *I. ricinus* vary from 0% to 66% [25], resulting in highly variable human exposure from tick bites. The reason for this high variability is unknown and could be attributed to the capacity of *R. helvetica* to be beneficial for the tick host under certain environmental conditions [31] or it could be related to genetic variation within *R. helvetica* populations, but the latter has not been investigated so far.

The pathogenic potential of *R. helvetica* has been explored, to a certain extent, on the cellular level. While other rickettsial species, which are pathogenic to humans, utilise actin-based cell-to-cell motility [32], a culture of *R. helvetica* has been shown to lack this ability in vitro [24], possibly due to a deletion in the *sca*2 gene, encoding a protein required for virulence that stimulates actin filament assembly [33, 34]. In addition, although the sequence encoding the RickA protein, a bacterial actin nucleator [35], was complete, its expression was apparently insufficient to enable actin-based motility in the strains studied [24]. Comparative analyses of its genetic makeup and metabolic potential are needed to clarify the pathogenic potential of *R. helvetica* as well as its interaction and localization within the tick host.

Advances in sequencing technologies have ushered in a new era of understanding by enabling the comprehensive annotation of complete tick mitochondrial genomes (mt genomes). These genomic insights have significantly enriched our understanding of tick phylogeny, population genetics, and evolutionary processes [36, 37]. Given ticks’ intimate association with symbiotic microbes crucial for their hematophagous lifestyle [38], the incorporation of mt genomes into co-phylogeny analyses presents a promising avenue for unravelling the evolutionary patterns of tick symbionts [39]. This approach holds particular significance for symbionts like *M. mitochondrii* which inhabit tick mitochondria [40] offering a unique window into their evolutionary trajectory.

Genome-scale model (GEM) reconstruction and analysis can further enhance this understanding by detailing the metabolic interactions and dependencies between ticks and their symbionts. These models can elucidate how these endosymbionts may contribute to the metabolic network, revealing potential targets for controlling tick populations and preventing tick-borne diseases. Additionally, GEMs can provide insights into the co-evolutionary dynamics of ticks and their symbionts, as has been shown in recent works [41], offering a clearer picture of how these complex biological systems have adapted and evolved together over time.

In this study, we applied deep sequencing on sixteen *Ixodes ricinus* specimens to unveil the genetic variation and differences in transmission dynamics of *M. mitochondrii* and *R. helvetica*. Complete genomes were obtained directly from the individual ticks using a hybrid assembly approach combining PromethION and Illumina sequencing. In addition to interspecies whole genome comparisons, we determined the genetic association of *R. helvetica* and *M. mitochondrii* with their tick hosts to draw conclusions about the mode of transmission of these symbionts. To further our understanding of the latter, we visualised *R. helvetica* in tick organs.

## 1 Methods

### 1.1 Tick specimen metadata, DNA isolation and sequencing and, Data FAIRification

#### 1.1.1 Tick specimen metadata and processing

Field samples: Six questing I. ricinus adult females, Ir_f1, Ir_f3, Ir_f6, Ir_f11, Ir_f12, Ir_f16. were collected by blanket dragging in a planted forest region (f) in the Netherlands. Another eight adult females were collected in a coastal sand dune coastal area (d) in the Netherlands and labelled as Ir_d1, Ir_d2, Ir_d3, Ir_d4, Ir_d5, Ir_d6, Ir_d7, Ir_d9. Exact locations are in (Table 1). Ticks were kept at −80°C until further processing Lab-reared: Questing I. ricinus nymphs were collected by blanket dragging and morphologically identified to species level using an identification key [42]. Nymphs were fed on an artificial membrane blood feeding system, as described in [43]. Two lab reared adults, (F1 and F2), were used in this study.

**Table 1.**
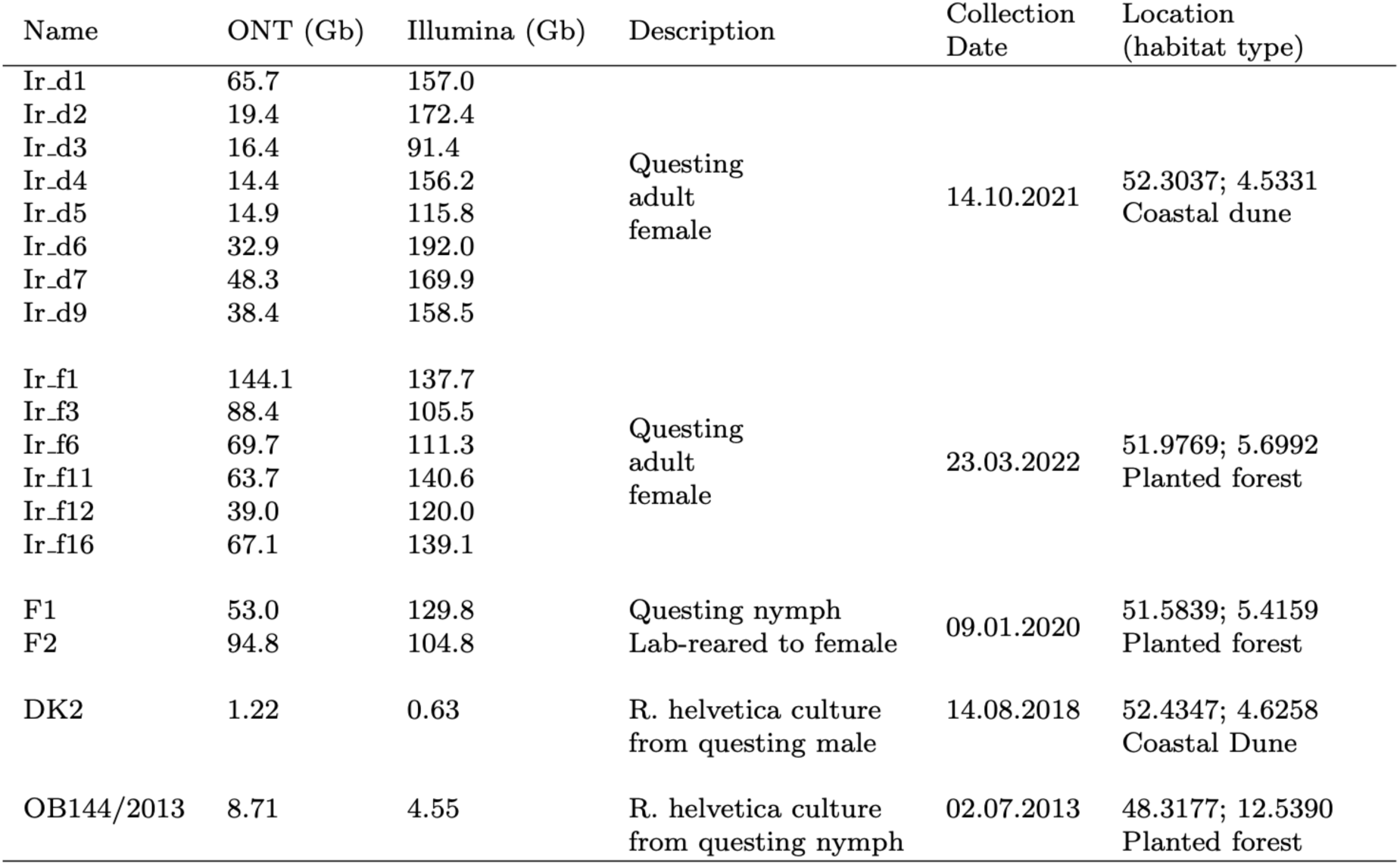
Origin and deep sequencing of adult *Ixodes ricinus* ticks.

#### 1.1.2 DNA isolation and sequencing of biospecimens

DNA extraction and sequencing was performed at Future Genomics Technologies BV (Leiden, The Netherlands). Ticks were ground to a fine powder using a pestle and mortar with liquid nitrogen. To extract high molecular weight DNA, a Genomic-tip 20/G kit (Qiagen Benelux BV, Venlo, The Netherlands) was used as per protocol.

DNA quality was measured via electrophoresis in Genomic DNA ScreenTape on an Agilent 4200 TapeStation System (Agilent Technologies Netherlands BV, Amstelveen, The Netherlands) and total DNA using a Qubit 3.0 Fluorometer (Life Technologies Europe BV, Bleiswijk, The Netherlands).

One microgram of the DNA samples was used to prepare a 1D ligation library using the Ligation Sequencing Kit (SQK-LSK110) as per protocol. (Oxford Nanopore Technologies (ONT), Oxford, United Kingdom). ONT libraries were tested on a MinION flowcell (FLO-MIN106) and subsequently run on a PromethION flowcell (FLO-PRO002) using the following settings: basecall model: high-accuracy; basecaller version: Guppy 5.0.17. Parallel aliquots of the DNA samples used for ONT sequencing were used to prepare Illumina libraries using the Nextera DNA Flex Library Prep Kit as per protocol (Illumina Inc., San Diego, CA, USA). Library quality was measured using an Agilent 4200 TapeStation System as described above. Genomic paired-end (PE) 150 nt libraries were sequenced using an Illumina NovaSeq 6000 system.

#### 1.1.3 DNA isolation and sequencing of R. helvetica strains from Vero cells

*Rickettsia helvetica* strain OB144 originated from a questing nymph and DK2 from a questing adult male tick. (Table 1). Strains were cultured and harvested from VERO cells as described [44, 45] with two modifications: here the needle and syringe protocol was implemented using 26 gauge needles and a 0.45µm syringe-driven membrane filter. Genomic DNA was extracted using the ZymoBIOMICS™ 96 MagBead DNA Kit (Zymo Research, Orange, CA). OB144 was sequenced by Future Genomics Technologies BV as described above. DK2 was sequenced by Baseclear (Leiden, the Netherlands) using a NovaSeq 6000 system (Illumina, San Diego, CA, USA). DK2 long reads were generated in house using a Nanopore GridION sequencer.

#### 1.1.4 Data FAIRification

The FAIR Data station [46] was used to register and submit the raw data and metadata associated with this study. The ENA “host-associated” checklist (ERC000013) was selected to meet the minimum metadata requirements for describing the biological samples.

### 1.2 Genome assemblies and annotation

A standardized hybrid workflow for metagenomics assembly [47] was used, which is registered at the WorkflowHub [48]. llumina and nanopore reads were inspected for quality using FastQC v0.12.1 [49]. Illumina reads were filtered with fastp v0.23.2 [50] with parameters –correction, –dedup, –disable trim poly g –length required 50, –qualified quality phred 20, –unqualified percent limit 20. Possible Illumina spike in PhiX genomic reads contamination was filtered with BBMap/bbduk v38.95 [51] -k 31, reference (NC 001422.1). To filter illumina reads for selected species BBMap v38.95 [51] was used with default settings and bloom=t. Nanopore reads were filtering using minimap2 v2.24 [52] default setting, –x and map-ont. Mapped reads were kept. Genome sequences used for filtering were extracted from the European Nucleotide Archive.

For *Rickettsia* all available *Rickettsia* reference genomes were used. For *Midichloria mitochondrii* the reference genome was (CP002130.1). For mitochondrial reads, all available *Ixodes* mitochondrial genomes were used. A lists of genome accessions for filtering can be found in supplementary File 1.

### 1.3 *R. helvetica* rickA gene variability analyses

Questing nymphs were collected in Duin & Kruidberg, Schiermonnikoog, Gieten, Veldhoven, Montferland and Wassenaar, the Netherlands as described [53], immersed in 70% alcohol and stored at −20°C. DNA was extracted by alkaline lysis as described [54]. Detection of R. helvetica *rickA* was done in two steps: qPCR of *gltA* gene followed by a rickA PCR assay targeting a 600 nt fragment of the rickA gene on qPCR positive samples, as described [55]. DNA-free water was used as negative control. Primers used: RIC1200 F2 GGCAAAATGTTAAAAATGTTT and RIC+430 R3 CCRGYTTTTTAACCGTAGTAG. RickA PCR products and the *rickA* gene sequences from the new R. helvetica genomes were aligned with ClustalW using Geneious 11.1.4 software [56].

### 1.4 Phylogenetic analysis of *Ixodes ricinus* mitogenomic data

ClustalW alignments were computed using Geneious 11.1.4 software [56]. Phylogeny was inferred by IQTREE 2.1.3 [57], and the best-fit evolution model was selected based on the Bayesian information criterion (BIC) computed by implemented ModelFinder [58]. Branch supports were assessed by the ultrafast bootstrap (UFBoot) approximation [59] and the Shimodaira–Hasegawa like approximate likelihood ratio test (SH-aLRT) [60]. Tree was rooted at the midpoint. No out-group was included. Tree was visualized and edited in FigTree (v1.4.1) and Inkscape (v 0.91) [61].

### 1.5 Cophylogenetic analysis

The single-copy core genes of 11 *R. helvetica* and 12 *M. mitochondrii* genomes were identified in Anvi’o 7 and made up of 1484 and 1127 single-copy gene (SCG) clusters, respectively. Gene alignments from SCG were extracted and concatenated using the program anvi-get-sequences-for-gene-clusters, with parameters: – concatenategene-cluster; –report-DNA-sequences. Maximum likelihood phylogenetic trees were constructed in IQ-TREE v2.2.0 [62]. The best model according to the Bayesian Information Criterion (BIC) was selected with Model Finder [63] as implemented in IQ-TREE. All models were run with Ultrafast Bootstrapping [64]. eMPRess, a program for phylogenetic tree reconciliation under the duplication-transfer-loss (DTL) model, was used to determine whether the mitochondrial genome of *I. ricinus* and *I. ricinus* symbionts: *M. mitochondrii* and *R. helvetica* coevolved by comparing their phylogenetic relationships. The input trees were based on the results of phylogenetic analyses. The analysis was conducted using the default eMPRess parameters.

### 1.6 Genome-Scale Metabolic Model Reconstruction and Analysis

Genome-scale metabolic models (GEMs) were developed using CarveMe (v1.5.1) [65], starting with their annotated genome sequences (Ir_d9Mm and Ir_d9Rh). For Reaction Essentiality Analysis, we performed gene or reaction deletion analysis using flux balance analysis (FBA). (Supplementary File 2). MEMOTE (v0.13.0) [66] was used to assess the quality of the GEMs. COBRApy (v0.26.3) [67] employing Python (v3.11.3), was used for modelling simulation and analysis of each GEM using the IBM CPLEX Solver.

### 1.7 Immunofluorescence localization studies of *R. helvetica* in ticks

Fifteen *I. ricinus* questing female ticks collected in the Amsterdamse Waterleidingduinen, the Netherlands and morphologically identified to species level using an identification key [42]. All ticks were kept alive until further processing. Ticks were dissected, and gut, malpighian tubules, salivary glands, reproductive system and synganglion were collected. Tick carcasses were screened for the presence of *R. helvetica* DNA by qPCR. The same was performed on 10 control ticks from a laboratory colony (BIPAR, France).

Organs were fixated in 4% paraformaldehyde (PFA) in PBS for 2h and then replaced by 70% ethanol and stored at −20C until used. The organs were washed by 0.5% Triton X-100 in PBS (PBST) five times, then replaced by 5% normal goat serum (Sigma) in PBST and incubated for 1h in room temperature. Afterwards, the serum was replaced by *R. conorii* commercial antibody 1:30 in PBST and incubated at 4C for 2 days. Samples were incubated with DAPI before mounting in Prolong Antifade Diamond Mountant (Life Technologies) and images were obtained using a z-stack tool in Leica DMI8 confocal microscopy. Assembled z-stacks were adjusted in Adobe Photoshop version 24.0.1 (Adobe Inc.).

### 1.8 Statistical analysis

Haplotype diversity (Hd), variance of Hd, nucleotide diversity (*⇡*), variable (polymorphic) sites, and parsimony informative sites were calculated for mitochondrial genomes of *I. ricinus* using the DNA SP6 (v6.12.03) [68]. To test whether the composition of repeats differs per prevalence and per area type (dune vs forest), we performed PERMANOVA based on Bray-Curtis dissimilarity matrix using adonis2 function from the vegan package [69].

## 2 Results

### 2.1 Deep sequencing of *I. ricinus* adult females reveals the presence and diversity of symbiotic bacteria

Sixteen adult female specimens were sequenced using both Illumina and ONT (Oxford Nanopore Technologies) methodologies: eight from a coastal sand dune area (d), six from a planted forest area and two specimens moulted under laboratory conditions. Using Illumina 150 x 150 nt paired-end reads as input and the NT database [70] as a reference, we applied a Kraken2-Bracken workflow [71,72] to estimate for each specimen, the abundances and taxonomic classifications of the microbiome-related reads obtained (Figure 1). In total, more than 90% of the read-pairs that were attributed to the bacterial domain could be classified at the species level.

**Fig. 1.**
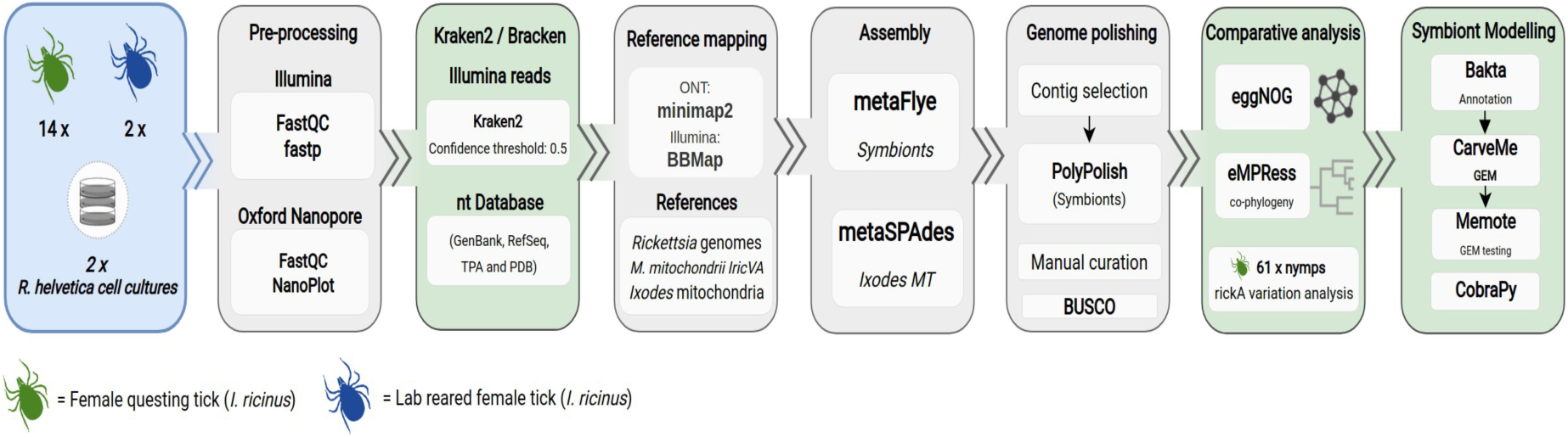
Ixodes ricinus endosymbiont genome assembly and analysis. A standardized workflow for hybrid metagenomics assembly was used (see Methods section for details). Two sources of genomic DNA were used as input; *I. ricinus* metagenomic DNA obtained from adult female ticks and *R. helvetica* genomic DNA isolated from infected Vero cell lines.

A significant number of paired-end reads (with cumulative scores of *>*20.000 read pairs) were assigned to known tick borne pathogens: *Anaplasma phagocytophilum* was present in tick Ir_d1 and Ir_d9; *Borrelia afzelii* was present in tick Ir_d9 and *Neoehrlichia mikurensis* [73] in tick Ir_f3. *Rickettsiella* species have recently been identified as facultative endosymbionts of *I. ricinus* ticks [74] and with more than 1 million read pairs assigned, *Ricketssiella* species appear to be present in Ir_d1, Ir_d6 and Ir_d7 (Figure 2).

**Fig. 2.**
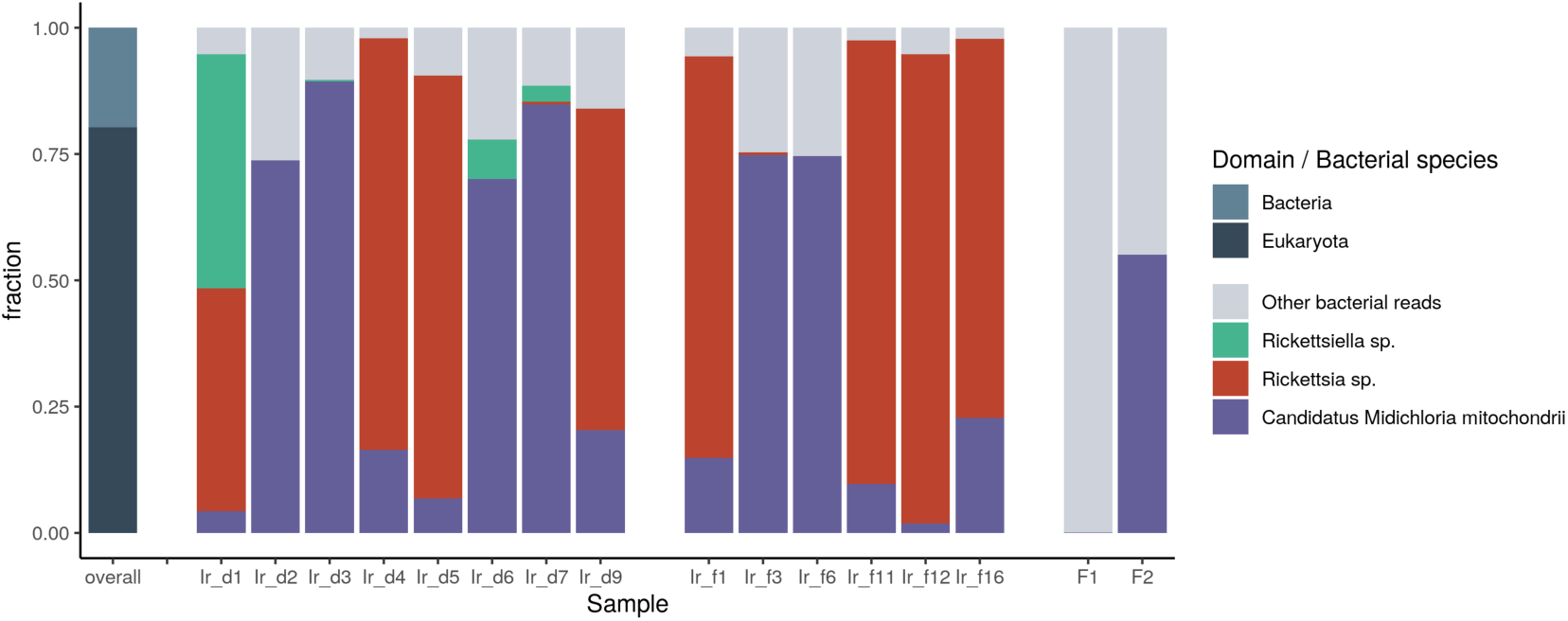
Bracken distribution of classified Illumina paired-end reads at the species level. Default parameters for kmer length (35), minimizer length (31), and minimizer spacing (7), were used in this analysis. Left: top level classification, right: classification of the bacterial domain at species level. The top 3 most prevalent species are indicated. The full analysis is available in Supplementary File 1.

Of the 14 specimens collected in the field (Table 1), eight tested positive for the presence of *R. helvetica* DNA by qPCR analysis (Supplementary File 1). Kraken analysis using the complete NT database as reference however, suggested that *Rickettsia asiatica*, *Rickettsia conorii*, and *R. helvetica*, three species that share almost identical genome sequences, appeared to exhibit a significant concurrent presence in these ticks. We used fastANI [75] to gain more insight into their evolutionary distances. A commonly used operational definition for prokaryotes states that if two genomes have an average nucleotide identity (ANI) of over 95%, they belong to the same species. Between them the reference strains of *R. helvetica* and *R. asiatica* show *>*98.66% ANI and in cross comparison with R. conorii *>*93.3% ANI. While these high identity scores explain the Kraken results, it’s important to note that FastANI revealed significant structural variation (Figure 3).

**Fig. 3.**
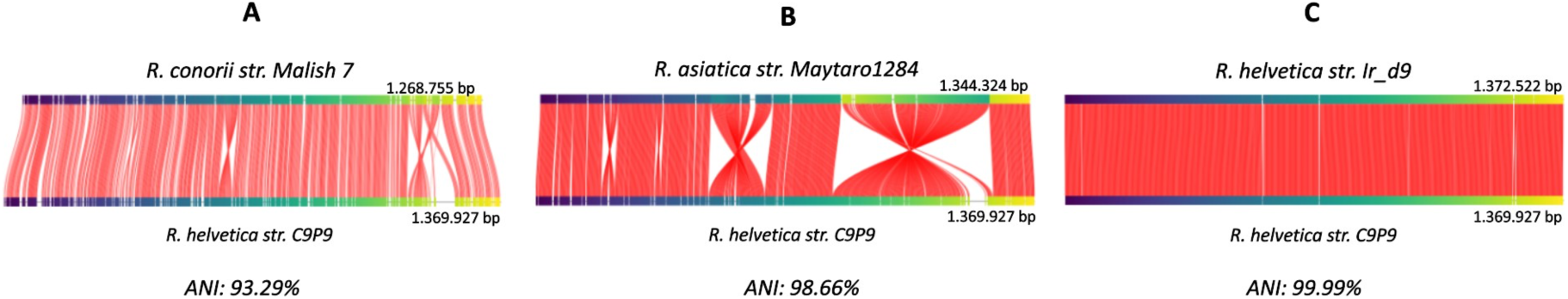
Major intragenomic rearrangements between *Rickettsia helvetica* and *Rickettsia asiatica.* FastANI orthologous mappings was done using a 3kb fragment size. Red line segments denote the orthologous mappings between these fragments. Panel **A**: ANI computed between *R. conorii* strain (NC 003103.1) as query and *R. helvetica* strain C9P9 (NZ CM001467.1) as reference. **B**: ANI computed between *R. asiatica* strain (NZ AP019563) as query and *R. helvetica* strain C9P9 as reference. **C**: ANI computed between the *de novo* assembled *Rickettsia* strain from tick Ir_dr9 as query and *R. helvetica* strain C9P9 as reference.

### 2.2 Reconstruction and phylogenetic analysis of the *I. ricinus*

#### mitochondrial genomes

MT genome size ranged from 14562 to 14575 bp. For all 16, a circular genome was obtained coding for 13 protein encoding genes, 22 transfer RNA genes (tRNAs) and 2 ribosomal RNA genes (rRNAs) (Supplementary File 1). The AT content of the MT genomes ranged from 79.2% to 79.3%, a characteristic feature shared with other hard tick species [76].

A multiple alignment of the 16 MT sequences indicated 214 variable positions of which 147 were informative parsimony sites (excluding indels) and a mean nucleotide diversity of 0.00460. These variable sites defined a total of 15 haplotypes (mean haplotype diversity, Hd = 0.9917, with samples Ir_f1 and Ir_f3 carrying the same haplotype. The phylogenetic analysis of the mitogenomic data suggests that a locality-dependent structure based on geographic location (dune and forest) is absent (supplementary figure 2).

### 2.3 Genome reconstruction of *M. mitochondrii* endosymbiotic strains

Kraken analysis revealed that 15 ticks are positive for *M. mitochondria*. The exception was F1, raised in a laboratory setting (Figure 2). High quality MAGs of *M. mitochondrii* could be obtained from 11 ticks and assembly drafts from the other four (Supplementary File 1). The MAGs shared over 99% ANI with the *M. mitochondrii* reference strain [77] and were characterised by high a BUSCO score (*>*92.9%). Genome sizes ranged from 1.190.312 to 1.191.611 bp with the exception of the assembly obtained from tick Ir_d2 which was 1.268.036 bp due to a duplication event (supplementary file 1). The average guanine cytosine (GC) content was 36.6%. All *M. mitochondrii* genomes contained 3 rRNAs, between 35-39 tRNAs, and the number of protein encoding genes varied from 1322 to 1448.

Among the eleven *de novo* Midichloria genome assemblies, a high level of genomic collinearity was observed (Figure 3, panel B, supplementary file 1) however, in comparison with the reference strain, a significant amount of structural variation was observed (supplementary file 2). Genome variant calling revealed strain variations. The total number of variants ranged from 604 to 964 (8509 total). Of these, 5029 single nucleotide polymorphisms (SNPs), 2246 insertions, 1989 deletions, and 335 complex variants were detected (Table 2).

**Table 2.**
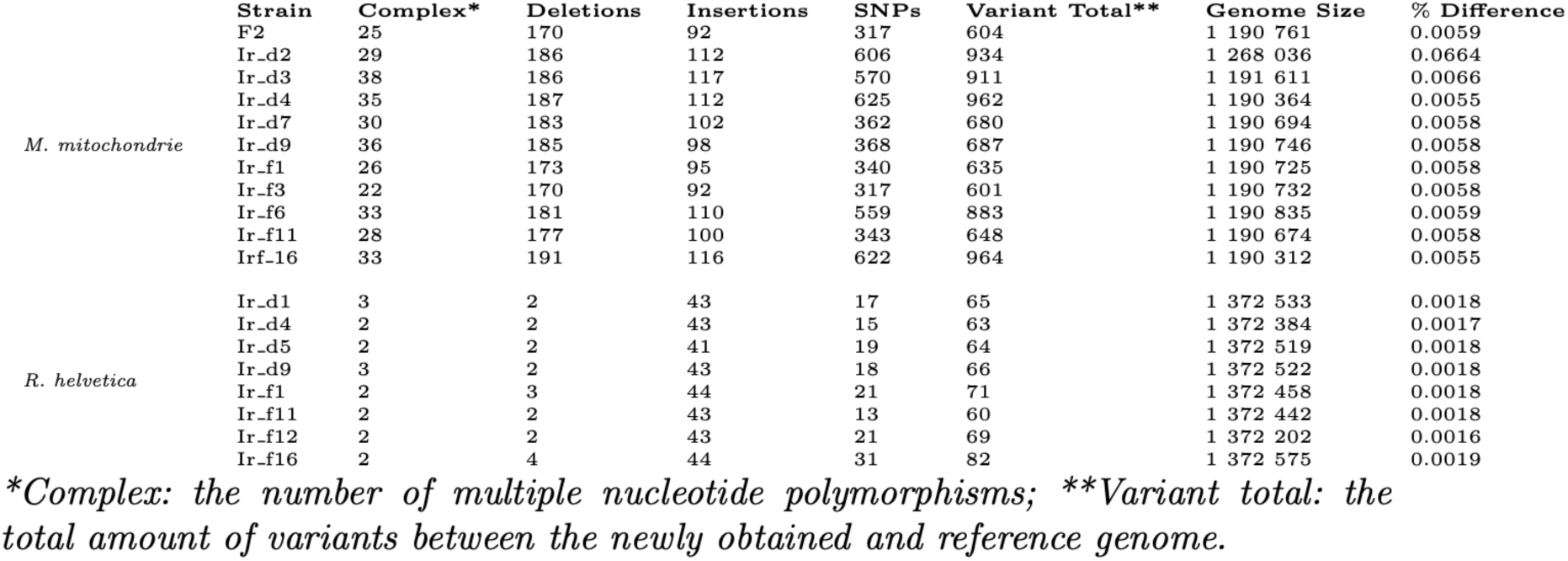
Genetic variation for 11 genomes of *Midichloria mitochonodrii* and 8 genomes of *Rickettsia helvetica* obtained in this study.

|  | Strain | Complex* | Deletions | Insertions | SNPs | Variant Total** | Genome Size | % Difference |
| --- | --- | --- | --- | --- | --- | --- | --- | --- |
| *M. mitochondrii* | F2 | 25 | 170 | 92 | 317 | 604 | 1 190 761 | 0.0059 |
|  | Ir_d2 | 29 | 186 | 112 | 606 | 934 | 1 268 036 | 0.0664 |
|  | Ir_d3 | 38 | 186 | 117 | 570 | 911 | 1 191 611 | 0.0066 |
|  | Ir_d4 | 35 | 187 | 112 | 625 | 962 | 1 190 364 | 0.0055 |
|  | Ir_d7 | 30 | 183 | 102 | 362 | 680 | 1 190 694 | 0.0058 |
|  | Ir_d9 | 36 | 185 | 98 | 368 | 687 | 1 190 746 | 0.0058 |
|  | Ir_f1 | 26 | 173 | 95 | 340 | 635 | 1 190 725 | 0.0058 |
|  | Ir_f3 | 22 | 170 | 92 | 317 | 601 | 1 190 732 | 0.0058 |
|  | Ir_f6 | 33 | 181 | 110 | 559 | 883 | 1 190 835 | 0.0059 |
|  | Ir_f11 | 28 | 177 | 100 | 343 | 648 | 1 190 674 | 0.0058 |
|  | Ir_f16 | 33 | 191 | 116 | 622 | 964 | 1 190 312 | 0.0055 |
| *R. helvetica* | Ir_d1 | 3 | 2 | 43 | 17 | 65 | 1 372 533 | 0.0018 |
|  | Ir_d4 | 2 | 2 | 43 | 15 | 63 | 1 372 384 | 0.0017 |
|  | Ir_d5 | 2 | 2 | 41 | 19 | 64 | 1 372 519 | 0.0018 |
|  | Ir_d9 | 3 | 2 | 43 | 18 | 66 | 1 372 522 | 0.0018 |
|  | Ir_f1 | 2 | 3 | 44 | 21 | 71 | 1 372 458 | 0.0018 |
|  | Ir_f11 | 2 | 2 | 43 | 13 | 60 | 1 372 442 | 0.0018 |
|  | Ir_f12 | 2 | 2 | 43 | 21 | 69 | 1 372 202 | 0.0016 |
|  | Ir_f16 | 2 | 4 | 44 | 31 | 82 | 1 372 575 | 0.0019 |

\*Complex: the number of multiple nucleotide polymorphisms; \*\*Variant total: the total amount of variants between the newly obtained and reference genome.

EggNOG based functional analysis suggests that the reference genome and the eleven newly sequenced genomes encode a (near) complete Folate biosynthetic pathway (6 of the 7 required genes detected) and a likely complete Biotin biosynthetic pathway (all required gene detected). COG0331 is present in all newly sequenced genomes but absent from the reference genome. COG0331 corresponds to malonyl CoA-acyl carrier protein transacylase, encoded by the fadD gene, and is responsible for initiating the type II fatty acid synthesis pathway [78]. A complete eggNOG based functional analysis is provided in Supplementary File 1.

### 2.4 Genome reconstruction of *R. helvetica* endosymbiotic strains

Of the 14 specimens collected in the field (Table 1), eight tested positive for the presence of *Rickettsia* strains by both qPCR and Kraken2 analysis and we were able to obtain the respective genomes and associated plasmids for all of them. Two additional genomes (DK2 and OB144) were obtained using of Vero cell lines.

All ten *de novo* assemblies were characterised by a high BUSCO score (*>*98.9%). Genome sizes varied from 1.372.202 bp to 1.372.575 bp which is about 2,2 2.5 kb larger than the C9P9 reference genome (1.369.927 bp). Consistent with the reference strain, all assemblies included a plasmid of 47184 48139 bp encoding 56 to 59 proteins, including 39-40 hypothetical proteins. All new *R. helvetica* genomes contained 3 rRNAs, 33 tRNAs and the number of protein genes ranged from 1630 to 1664 per assembly including a complete set of genes for *de novo* folate biosynthesis (Supplementary File 1).

Genome variant calling among the eight *I. ricinus* derived genome assemblies revealed a total number of variants ranging from 63 to 82 (540 in total). Of these, 155 single nucleotide polymorphisms (SNPs), 344 insertions, 19 deletions, and 18 complex variants were detected (Table 2).

Overall the new (meta)genome assemblies revealed a very high level of genomic collinearity, among themselves, and with the *R. helvetica* reference strain (Figure 3 panel C, supplementary file 1). Most variation is due to what appears to be gene splitting events leading to two or more open reading frames. Between the eight genome assemblies that were directly obtained from the *I. ricinus* specimens we found three such gene splitting events. In contrast, we observed a notable increase of such events in the genomes of the DK2 strain and the C9P9 reference strain both obtained using infected Vero cells (supplementary File 1).

Due to such gene splitting event the DK2 genome encodes a non-functional RickA protein (Supplementary File 1). RickA plays a role in actin-based cell-to-cell motility and *rick* A is therefore considered to be a putative virulence gene. [32, 79]. Further inspection of the new *R. helvetica* genomes revealed high genetic variability in the rickA gene due to copy number variation of a 33-nt-long repeat motif encoding 11 amino acids including 5 adjacent prolines ranging in number from 5 (Ir_d4) to 13 (OB144) (supplementary File 1).

To gain more insight into this genetic variability, we sequenced the fragment of the *rick* A gene containing the repeat motif from 61 questing nymphs gathered from various forest and dune regions (supplementary File 1). The results confirmed the observed copy number variation which varied from 5 to 11. No significant compositional difference was observed between dune and forest areas (F 1,7 = 1.2, p = .3772), as well as between eight sampled locations with varying *R. helvetica* prevalence (F 1,7 = 0.4, p = .9101).

### 2.5 Genome-scale metabolic modelling of *M. mitochondrii* and *R. helvetica* reveal metabolic dependencies

Due to their adaptation to rely on the host organism for survival *M. mitochondrii* and *R. helvetica* have reduced genomes encoding reduced metabolic networks. The automatically generated Genome-scale metabolic model (GEM) of *M. mitochondrii* (Ir_d9 Mm) is comprised of 1126 reactions, 881 metabolites, and 266 genes, while the GEM of *R. helvetica* (Ir_d9 Rh) consists of 1123 reactions, 851 metabolites, and 274 genes (Table 3).

**Table 3.** Summary of genome-scale metabolic models of *M. mitochonodrii* and *R. helvetica*.

|  |  | Ir_d9_Mm | Ir_d9_Rh |
| --- | --- | --- | --- |
| Total No. of Reactions |  | 1126 | 1123 |
| Total No. of Metabolites |  | 881 | 851 |
| Total No. of Genes |  | 266 | 274 |
| Reactions | Transport Reactions | 349 | 399 |
|  | Metabolic Reactions | 630 | 581 |
|  | Without GPR associations | 337 | 353 |
| Metabolites | Unique metabolites | 881 | 851 |
|  | Medium components | 144 | 139 |
| Genes | Enzyme Complexes | 36 | 49 |

MEMOTE analysis [66] revealed that both GEMs achieved high-quality overall scores of 84% (Ir_d9 Mm) and 68% (Ir_d9 Rh), respectively. (supplementary File 3). The network motifs of the two models point to differences in central carbon and fatty acid metabolisms. For instance, the central carbon metabolism appears to be more tightly connected to nitrogen metabolism in the Ir_d9 Rh GEM compared to Ir_d9 Mm GEM. Despite these differences, the two GEMs share 802 reactions. Each GEM also contained unique reactions and pathways: 300 for Ir_d9 Rh GEM and 429 for Ir_d9 Mm GEM. More specifically, the Ir_d9 Mm GEM contained unique reactions and pathways related to pantothenic acid (vitamin B5) and niacin (vitamin B3). Reaction essentiality analysis indicates that 8.70% and 8.17% of the reactions in *M. mitochondrii* and *R.helvetica*, respectively, are essential for *in silico* growth (Supplementary File 2) and suggests that reactions linked to lysine synthesis pathways are vital for the growth of *R. helvetica* but not for *M. mitochondrii* (Figure 4)

**Fig. 4.**
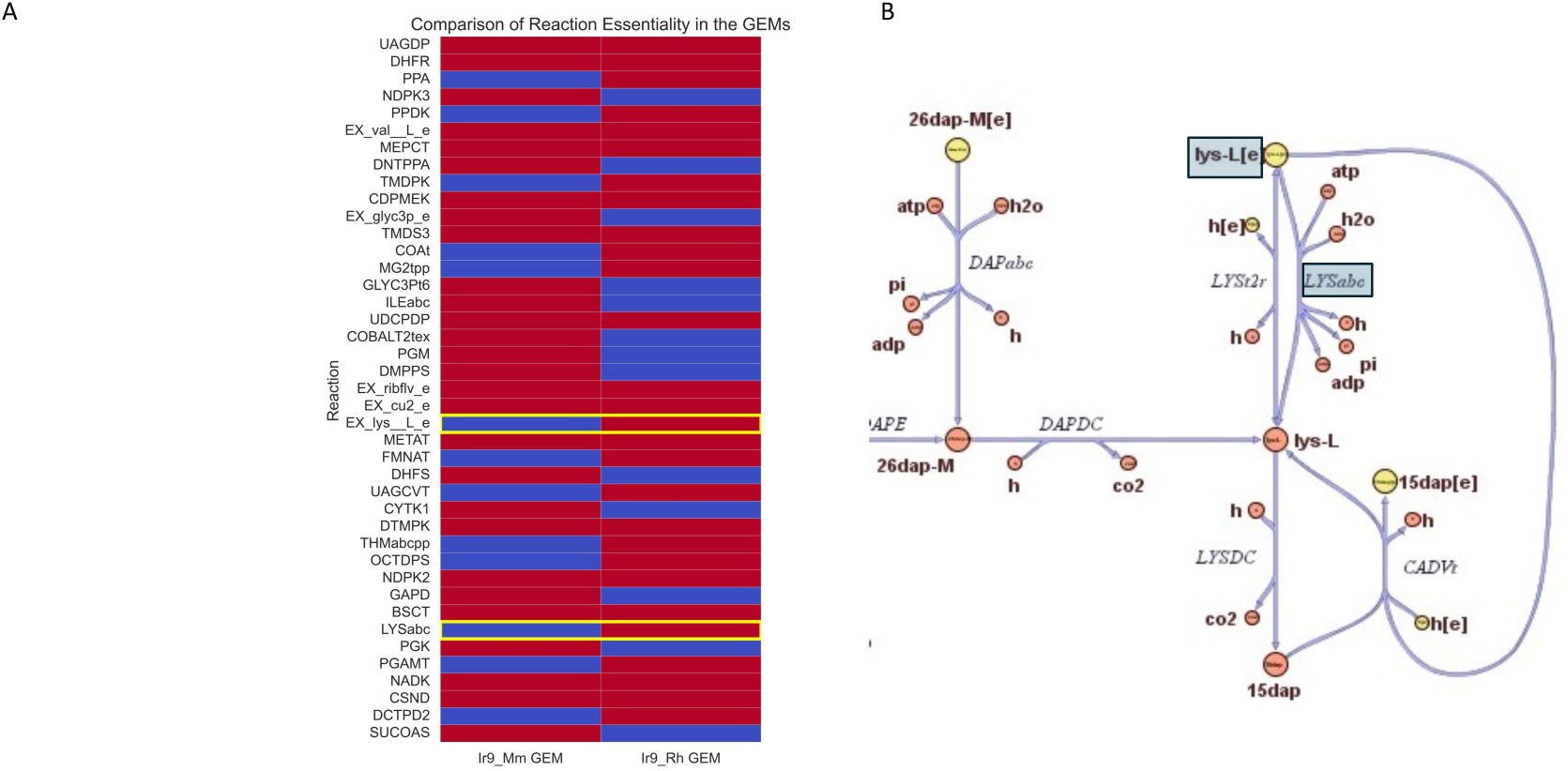
Genome-scale metabolic constructions and analysis of *M. mitochondrii* and *R. helvetica*. Heatmap comparing *M. mitochondrii* (Ir_d9 Mm GEM) and *R. helvetica* (Ir9 Rh GEM) where a subset of the essential reactions is depicted. Furthermore, blue and red panels represent which reactions are not essential or essential, respectively (panel A). Note: Complete heatmap comparing reaction essentiality of the two GEMs is contained in the supplementary material. Network map showing the network map of lysine synthesis pathway highlighting specific essential reactions in the Ir_d9 Mm GEM for *M. mitochondrii* (Ir_d9 Mm GEM) (panel B).

### 2.6 Cophylogenetic analysis between *I. ricinus* and symbionts

To determine whether symbiont-tick pairs are co-evolved, the eMPRess phylogenetic tree alignment software following the duplication-transfer-loss model was used to compare the phylogenetic relationships of symbionts and ticks. The analysis was performed separately for two pairs: *M. mitochondrii-I. ricinus* and *R. helvetica–I. ricinus* (Figure 5). The results showed that a total of 12 events, including eight cospeciation events, two transfers, and two losses, occurred between *M. mitochondrii* and *I. ricinus*, over evolutionary time; of these events, six were robustly supported. The results of this analysis refuted the null hypothesis that the tick and *M. mitochondrii* trees and tip associations are formed due to chance at 0.01 level in host-symbiont relationships (P value = 0.0099). Therefore, we concluded that these *I. ricinus M. mitochondrii* pairs have coevolved. The results for the *R. helvetica I. ricinus* pair did not indicate the rejection of the coevolution hypothesis (Pvalue = 0.77).

**Fig. 5.**
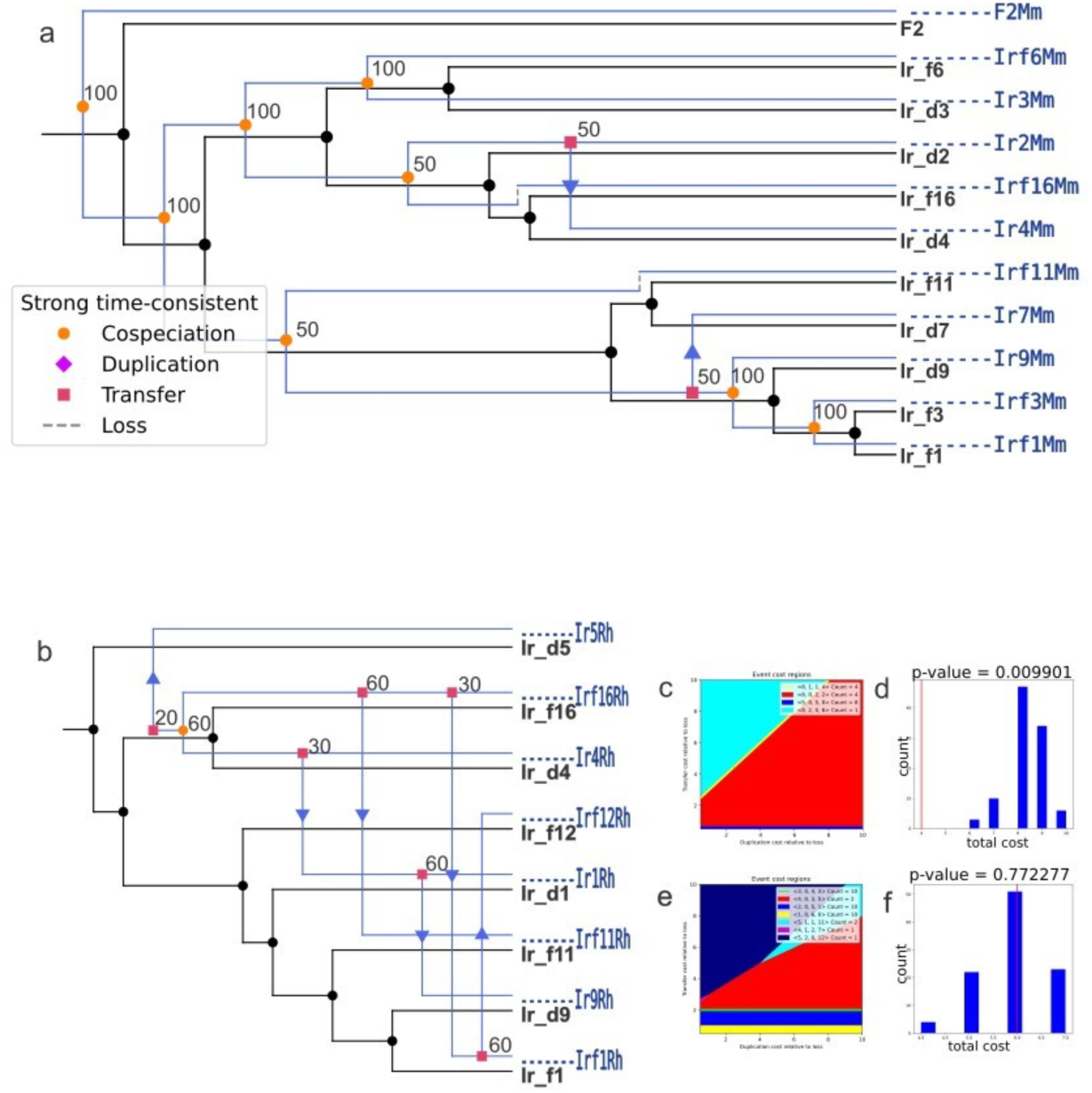
Cophylogeneticanalysisbetween *I. ricinus* ticks and its symbionts. a) Phylogenetic reconciliation of *I. ricinus* (displayed in black) and *M. mitochondrii* (in blue). b) Phylogenetic reconciliation of *I. ricinus* (displayed in black) and *R. helvetica* (in blue); c); d) P-value histogram of host and *M. mitochondrii* ; e); f) P-value histogram of host and *R. helvetica*; the optimal reconciliation cost of the coevolution trees is indicated with a red line, and the optimal cost of the same trees constructed with tip associations permuted at random is shown in blue columns.

### 2.7 *Rickettsia helvetica* localization in adult female ticks

The distribution of *R. helvetica* in adult female ticks was studied to examine the nature and uniformity of its presence in tick tissues and to identify possible transmission routes. *Rickettsia helvetica* was successfully visualised using laser microscopy in 5 out of the 15 females (Figure 6). These ticks were also qPCR positive for *R. helvetica*. All controls were negative (supplementary file 4). *Rickettsia helvetica* was observed in salivary glands, malpighian tubules, in the gut and in reproductive organs (supplementary file 4). The distribution of *R. helvetica* in tick organs varied between specimens (supplementary file 4).

**Fig. 6.**
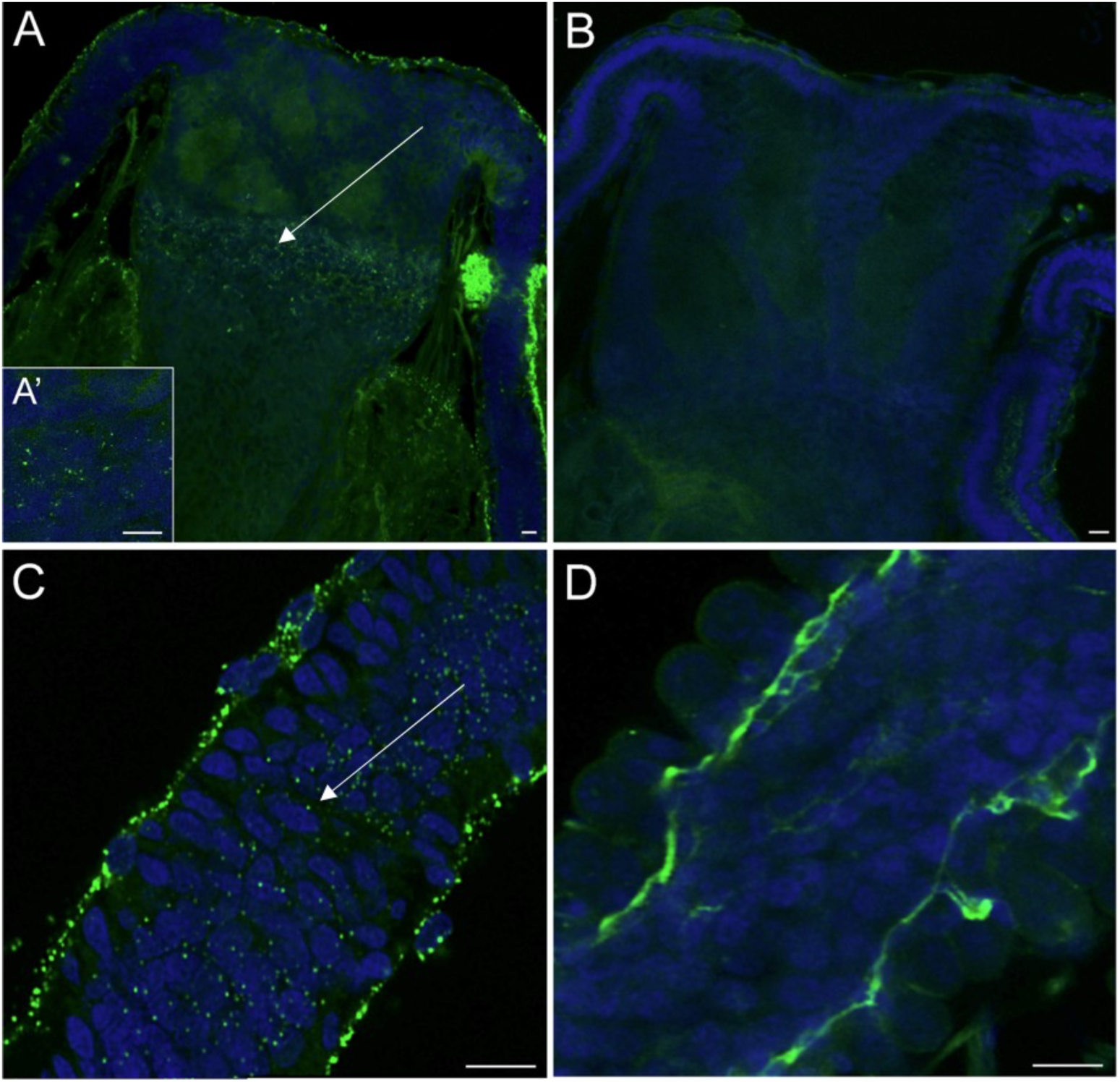
Localization of *Rickettsia helvetica* in adult females., Blue: DAPI, Green dots: *Rickettsia* specific binding. A) R. helvetica positive *receptaculum seminis* X40 A’) *R. helvetica* positive *receptaculum seminis* X100, B) Control *receptaculum seminis*, C) *R. helvetica* positive oviduct X40, D) Control oviduct. Note that tissue specific autofluorescence can be observed in the treatment and control groups. Scale-bar: 10*µ*m

In the reproductive system, *R. helvetica* was observed in the female oviduct (2 ticks, Figure 6 C,D) and in the *receptaculum seminis* (RS) (3 ticks, Figure 6 A,B), which acts as the storage site for sperm introduced during copulation.

## 3 Discussion

### 3.1 Endosymbiont genome sequencing

Conducting whole-genome sequencing of obligate intracellular bacteria often involves essential time and resource consuming steps [80]. These steps include isolating the microorganism from the host, infecting an intermediate replication host with the associated risk of host adaptation, and purifying the microbial DNA from this intermediate host. To address these challenges, we employed a direct deep-sequencing approach to acquire genomes of *M. mitochondrii* and *R. helvetica*, aiming to investigate their transmission dynamics and lifestyle.

The *I. ricinus* holobiome is still being discovered. The *I. ricinus* genome has been only recently been assembled [81] and the microbial component is underexplored. We used Kraken2 to profile the bacterial domain, utilizing the NT database as a reference, which includes (partially) non-redundant nucleotide sequences from all traditional public databases, but not the WGS draft genome assembly database. Kraken2 employs exact-match database queries of k-mers to classify sequences within a dataset down to species level. At this level, Kraken2 was able to classify 46 species with a cumulative presence of more than 1000 paired-end reads assigned across the 16 individual female ticks. This number corresponds with results from the analysis of the complete microbiome of 106 female *I. ricinus* ticks using 16S rRNA gene amplicon sequencing. [82]. At genus level these samples were dominated by *Candidatus Midichloria*, present in 99.5% of the samples, followed by *Pseudomonas, Methylobacterium, Sphingomonas,* and *Rickettsia*. *Rickettsiella* was present in 60.9% of these tick samples.

In the current study *M. mitochondrii* is present in 15 specimens. The exception is a specimen molted under laboratory conditions. *Rickettsia helvetica* was present in eight, while *Rickettsiella* species were present in three specimens. Furthermore, we did see a high overall presence of *Pseudomonas, Mycobacterium* and *Mycobacteroides* species. Note that the sensitivity of a Kraken2 analysis is significantly influenced by the type of genetic data available for these species in the NT database being 16S sequences, individual gene (fragments) and*\*or complete genomes implicating that Kraken read counts cannot directly be used to estimate relative species abundances.

As there are reference genomes available for *M. mitochondrii* and *R. helvetica*, employing fixed-length Illumina read-pairs as input allowed us to directly utilize the Kraken2 output for estimating whether there exists adequate short-read genome coverage to reconstruct the complete microbial genomes of interest. In alignment with the outcomes from Kraken2, we were able to obtain eleven complete and four draft genomes of *M. mitochondrii* and, coinciding with qPCR analysis results, eight complete *R. helvetica* genomes along with their associated plasmids.

### 3.2 Midichloria mitochondrii

The genomes of *M. mitochondrii* as presented in this study are approximately 5% larger and have a structural arrangement that differs from the published assembly (Acc. No.: NC 015722.). These differences are likely due to geographic separation (the Netherlands vs. Italy). The extraction of 11 complete *M. mitochondrii* genomes, each from a single tick, provides insight into the genomic potential and structural variability of this symbiotic bacterium in its tick host. *Midichloria mitochondrii* exhibits a unique feature to colonize multiple compartments in a tick body, including ovaries, salivary glands [83], Malpighian tubules, tracheae and guts [84]. In this context, each genome generated in our study represents a mixture of genomes from bacteria, inhabiting different ecological niches within the tick body. Subsequently, in the eMPRess reconciliation analysis, we showed a significant coevolutionary relationship between

*M. mitochondrii* and its tick host. This finding supports the notion of a long-standing symbiotic relationship between *M. mitochondrii* and *I. ricinus*, indicating that the vertical mode is this symbiont’s main mode of transmission. Also, the observed strong cophylogeny between *M. mitochondrii* and its host’s mitochondria is in opposition to a hypothesis that this symbiont undergoes horizontal transmission. Although *M. mitochondrii* has been shown to be localized in tracheae, salivary glands, and rostrum as well as it has been detected in the blood of experimentally infested rabbits, it is unlikely that it utilizes horizontal transmission to spread within the tick population under natural conditions.

### 3.3 Rickettsia helvetica

The prevalence of *R. helvetica* in questing ticks has been shown to greatly differ between dune and forest areas in the Netherlands [25, 31, 85]. Although our findings point to some heterogeneity in the rickA virulence gene, there seems to be no genetic basis for the differences in prevalence of *R. helvetica* in dune versus forest areas. Thus two remaining possibilities should be further explored. Firstly, the possibility of *R. helvetica* giving ticks in dune areas an adaptive advantage in relation with local abiotic conditions has not been fully explored. The role of *R. helvetica* in this case could be related to nutrient acquisition, waste elimination, or to physical characteristics. For example, *Rickettsia* has been shown to enhance tick motility in *Dermacentor variabilis* [86]. Secondly, the possibility of interactions with other symbionts within the tick host should be explored as *R. helvetica* has been shown to modulate the microbiome of *I. ricinus* [87] and it has been found to be negatively correlated with certain symbionts such as *Spiroplasma ixodetis* and *M. mitochondrii* [2, 31]. It has been hypothesised that perhaps the biological roles of *S. ixodetis* and *R. helvetica* overlap, making their simultaneous maintenance energetically superfluous for the tick host [31], but this theory has yet to be proven. Nor has the option of other bacterial interactions been fully researched.

Our comparative analyses showed near absolute genetic homogeneity among the generated *R. helvetica* genomes (99.96-100%). and only a slight deviation from the published strain C9P9. In the latter case, the observed differences in genome size could be due to different isolation and assembly methods, or even geographic origin. The congruous genetic build-up among the *R. helvetica* genomes was punctuated almost exclusively by the gene encoding for the surface protein RickA. The sequence of the *rick* A gene showed high genetic variation between the genomes generated in this study as well as between genes obtained from 61 additional ticks from the Netherlands. RickA has been shown to induce actin polymerization, and hence actin-based motility, in some *Rickettsia* species [35, 88]. However, a previous study found the RickA protein in *R. helvetica* isolate AS819 to be insufficient to promote actin-based motility *invitro*, which spread instead through cell break-down and bacterial release [24]. The inability of the studied isolate to spread through actin-based motility was explained by a disrupted *sca*2 gene, another virulence gene believed to have a more prominent role than the *rick* A gene in the cell to cell spread of *Rickettsias* [89]. Our genomes revealed no truncated sca2 sequences, however, the rickA gene greatly differed in length due to a repeat motif. We hypothesise these repeats may affect the functionality of the rickA protein within *R. helvetica*, giving rise to variation in virulence. This observation is supported by the dose-dependent manner in which rickA induced cell motility in *R. conorii* [35]. Moreover, when expressing a heterologous rickA gene, *R. bellii* moved at higher velocities than its wild-type counterparts [90]. If the diverse rickA sequence length translates into different motility-associated phenotypes, it could explain the seemingly sporadic pathogenicity we observe for *R. helvetica*, as well as its low and inconsistent prevalence rate in wildlife hosts. This heterogeneity could also explain why two different mice models showed no substantial rickettsemia following an infection challenge with *R. helvetica* [91] and should be taken into account when exploring the infection potential of *R. helvetica* strains.

To date, no wildlife reservoir has been found for *R. helvetica*. This has led to the assumption that *I. ricinus*, which transmits *R. helvetica* transovarially [92], acts as its main reservoir in nature [25]. This close tick-symbiont interaction was not reflected in our findings which showed the absence of co-phylogeny between *R. helvetica* and the mitochondrial genomes of the *I. ricinus*. The lack of co-evolution between *I. ricinus* and *R. helvetica* may be partly due to a mixed transmission strategy encompassing vertical and horizontal routes. The presence of *R. helvetica* in the RS, as shown by our immunofluorescence findings, strongly suggests the possibility of paternal transmission for *R. helvetica* in *I. ricinus*. Paternal transmission of *Rickettsia* has been identified in the leafhopper *Nephotettix cincticeps* [93] and it has been hypothesised that a mixed mode of transmission (maternal and parental) might allow for mixing of different symbiont lineages thus allowing for the evolution of virulent variants [94]. *Rickettsias* have also been detected in the spermatids of *Dermacentor silvarum* and *I. ricinus* ticks [95, 96]. Thus, the possibility of a different genetic makeup for *Rickettsia* populations found in reproductive tick organs should be considered in future studies.

### 3.4 Symbiont metabolic integration

Genome-Scale Metabolic Models (GEMs) provide a means to uncover new biological insights that go beyond what genomics can offer. A key application involves predicting the viability of an organism under specific conditions. This simulation approach has been utilized to identify potential metabolic drug targets, enabling the effective targeting of pathogens [97].

Due to their long-term relationship with the tick host, both *M. mitochondrii* and *R. helvetica* underwent genome reduction, leading to incomplete metabolic networks. The analysis suggests that *M. mitochondrii* ‘s *in silico* growth necessitates the inclusion of pantothenate (vitamin B5) and glycerol-3-phosphate (G3P) in the growth medium. G3P, a key metabolite in metabolic pathways such as glycolysis, gluconeogenesis, and lipid metabolism, is abundantly available in mitochondria. Its role is pivotal in the glycerol phosphate shuttle, facilitating the transfer of reducing equivalents to the mitochondrial matrix. Pantothenate is vital for energy metabolism and the synthesis of coenzyme A (CoA), a coenzyme integral to mitochondrial metabolic reactions. Our findings about the growth requirements of *M. mitochondrii* are in line with the evolutionary adaptation of this symbiont to reside in its host’s mitochondria. Nevertheless, as ticks cannot synthesize pantothenate, there could be potentially other bacterial symbionts producing this vitamin to be utilized by *M. mitochondrii*. The GEM for *R. helvetica* highlights a dependency on lysine for *in silico* growth, as demonstrated in (Figure 4B). This requirement likely results from an incomplete lysine biosynthesis pathway, a characteristic shared with *Rickettsia prowazekii* [98, 99], as previously identified. Research has consistently shown that *Rickettsia* species lack comprehensive amino acid biosynthesis pathways, particularly for essential amino acids such as lysine [100, 101]. Therefore, *R. helvetica* must obtain lysine from the blood meal ingested by its tick host or from other tick symbionts, however, the latter has not been explored. In some cases, *Rickettsia* species have been found to carry out only the initial stages of lysine biosynthesis, leading to the production of diaminopimelate rather than lysine itself, necessitating the acquisition of lysine from their external environment [100].

Simultaneously, we examined how these symbionts could contribute to their tick host by providing essential micronutrients. All *R. helvetica* genomes harbour a complete set of genes for de novo folate biosynthesis while in line with previous findings [102, 103]. *M. mitochondrii* encodes a (near) complete folate bio-synthetic pathway and a likely complete biotin biosynthetic pathway.

## 4 Conclusions

Ticks harbour and transmit specialized symbiotic microorganisms that are crucial for their hematophagous lifestyle and serve as causal agents of diseases in humans and animals. While these symbionts are challenging to culture and manipulate ex vivo, deep sequencing allows for the direct extraction of complete genomes from tick specimens collected from their natural habitats providing fresh insights into their structural genome variation, host-interactions and mode of transmission. The infectivity and transmission dynamics of R. helvetica may be influenced by genetic variations in the rickA virulence gene.

## 5 Data & Code availability

The data supporting the findings of this study have been deposited in the European Nucleotide Archive (ENA). This dataset includes Oxford Nanopore Technologies (ONT) and Illumina sequencing data for multiple *Ixodes ricinus* specimens.

|  |
| --- |
| **Project:**PRJEB70514 |
| **ONT:** ERR12319604,ERR12319605,ERR12319607,ERR12319608, ERR12319610,ERR12319612,ERR12319613,ERR12319614, ERR12319616,ERR12319618,ERR12319620,ERR12319623,ERR12319624,ERR12319627,ERR12319628, ERR12319631,ERR12319632,ERR12319593,ERR12319595,ERR12319597, ERR12319598,ERR12319599, ERR12319600, ERR12319602 |
| **Illumina:** ERR12319603, ERR12319606, ERR12319609, ERR12319611, ERR12319615,ERR12319617, ERR12319619, ERR12319621, ERR12319625, ERR12319626,ERR12319629, ERR12319630, ERR12319633, ERR12319622, ERR12319594,ERR12319596, ERR12319601 |

### Simulation code

The code used for the model simulations can be found at https://git.wur.nl/unlock/projects/ticks

### Genome-Scale Models

The GEMs as well as accompanying MEMOTE[66] and FROG Analysis[104] reports can be found in a public repository on BioModels: https://www.ebi.ac.uk/biomodels/. For the *M. mitochondrii* GEM, see here: MODEL2404170001. For the *R. helvetica* GEM, see here: MODEL2404170002.

## Declarations

B.N, W.T.S.J, P.J.S, and J.J.K acknowledge the Dutch Research Council (NWO), and Wageningen University & Research for their financial contribution to the Unlock initiative (NWO: 184.035.007). T.A. and H.S. were supported by funding from the Dutch Ministry of Health, Welfare and Sport.

## Supplementary information

**Supplementary File 1 (XLSX):** Information about the strains, coverage, busco, ani and tera analysis, genome accessions, eggNOG annotation, RickA and co-linearity analysis.

**Supplementary File 2 (PDF):** Genome-scale modelling details.

**Supplementary File 3 (PDF):** Phylogenetic tree of the mitochondrial *Ixodes ricinus*

**Supplementary File 4 (PDF):** *Rickettsia helvetica* localization in adult female ticks

**Code repository:** https://git.wur.nl/unlock/projects/ticks/-/tree/main/ supplementary material

